# Recovery of an 18^th^ Century Rhinovirus Genome through Ancient RNA Isolation of Human Lungs

**DOI:** 10.64898/2026.01.29.702071

**Authors:** Erin E. Barnett, Alejandra Castillo, Isabelle A. Du Plessis, Kathryn Kistler, Laura Carrillo, Anahí Sanchez Leon, Ting Liu, Mike G. Rutherford, Jon Ploug, John T. McCrone, María C. Ávila-Arcos, Daniel Blanco-Melo

## Abstract

RNA viruses cause substantial global morbidity, yet their impact prior to the twentieth century remains obscured. While ancient DNA studies have transformed our understanding of past pathogens, ancient RNA (aRNA) isolation is largely restricted to exceptionally preserved samples. Here, we simultaneously recover aDNA and aRNA from non–formalin-fixed human lung specimens and reconstructed an 18th-century Human Rhinovirus (HRV) A genome—the oldest human RNA virus identified to date. The RNA is highly fragmented, with distinctive terminal misincorporations and coverage patterns consistent with double-stranded RNA. Phylogenetic analyses indicate that this historical HRV genome is an extinct lineage related to contemporary genotypes, providing a unique perspective on rhinovirus evolution. These findings demonstrate that centuries-old medical specimens can retain informative aRNA, expanding the temporal scope of paleovirology.

## INTRODUCTION

Emerging and re-emerging RNA viruses have driven devastating global health crises in recent decades (*1*). Although historical accounts dating back to ∼1500 B.C.E. describe symptoms that could be attributed to viral infections (*2*), the impact of RNA viruses prior to the 20^th^ century remains largely unknown. Significant advances in paleogenomics have substantially improved our ability to characterize pathogens hidden within historical human remains (*3*–*6*). For instance, full-length reconstructions of numerous Hepatitis B viral genomes have illuminated the intricate co-evolution with their human hosts over the last 7,000 years (*7*–*9*). Despite this progress, recovering viral RNA genomes from historical specimens remains challenging due to the inherent limitations of ancient RNA (aRNA) isolation (*10*). RNA molecules largely exist as single stranded molecules and are biochemically less stable than double stranded RNA (dsRNA) and DNA (*11*). In addition, the release of RNases from tissues post-mortem and their ubiquitous presence in the environment further promote RNA degradation. As a result, aRNA isolation typically recovers fragments shorter than 30 nucleotides from specimens frozen in permafrost or desiccated tissues (*12*–*15*). Formalin-fixed tissues have shown promise for consistent aRNA recovery with less degradation than other historical sample types; and has resulted in the reconstruction of a limited number of viral RNA genomes, including Influenza A Virus and Measles Virus (*16*–*18*). Medical museums and biological archives provide unique opportunities to explore the history and evolution of RNA viruses of global health importance – however, formalin was not adopted as a preservative until early 1900s (*19*). Thus, leveraging these resources requires expanding the technical capabilities of aRNA isolation across alternative preservation methods. Here, we present the successful recovery of aRNA from non-formalin fixed tissue specimens dating to the 18^th^ and 19^th^ century, enabling the genome reconstruction of an 18^th^-century RNA virus – the oldest known human RNA virus genome to date.

## RESULTS

### Dual aDNA/aRNA isolation from human lung specimens associated with Great Britain’s Industrial Revolution

The rapid industrialization of Great Britain (c. 1760–1840 C.E.) was accompanied by a surge in infectious diseases, driven by poor living conditions and inadequate sanitation in densely populated areas (*20*). To investigate viral respiratory diseases in this historical context, we performed a detailed review of pathology records from The Hunterian Anatomy Museum (University of Glasgow, Scotland). Preserved in spirits of wine (alcohol) rather than formalin, wet specimens in the collection of Dr. William Hunter (1718–1783) represent a rare archive of pre-formalin biological material (**Supplementary Materials**). We identified two human lung specimens – GLAHM:120916 (c. 1740–1783) and GLAHM:120917 (c. 1848–1877) – with documented evidence of severe respiratory disease (**Fig. 1A & Supplementary Materials**). We successfully co-isolated DNA and RNA from both specimens using the Qiagen DNeasy Blood and Tissue Kit with key modifications for small nucleic acid recovery (**Fig. 1B & Supplementary Materials**). There were neglectable levels of nucleic acids in our environmental (preservation liquid) and negative (water) controls (**Fig. 1B**). High-throughput sequencing shows significant degradation of both RNA and DNA, with mean RNA fragment lengths of 28.05 and 24.40 nucleotides (nt) for samples GLAHM:120916 and GLAHM:120917, respectively, nearly half the average size of DNA fragments (average 48.99nt) (**Fig. 1C)**(*21*).

**Fig. 1.**
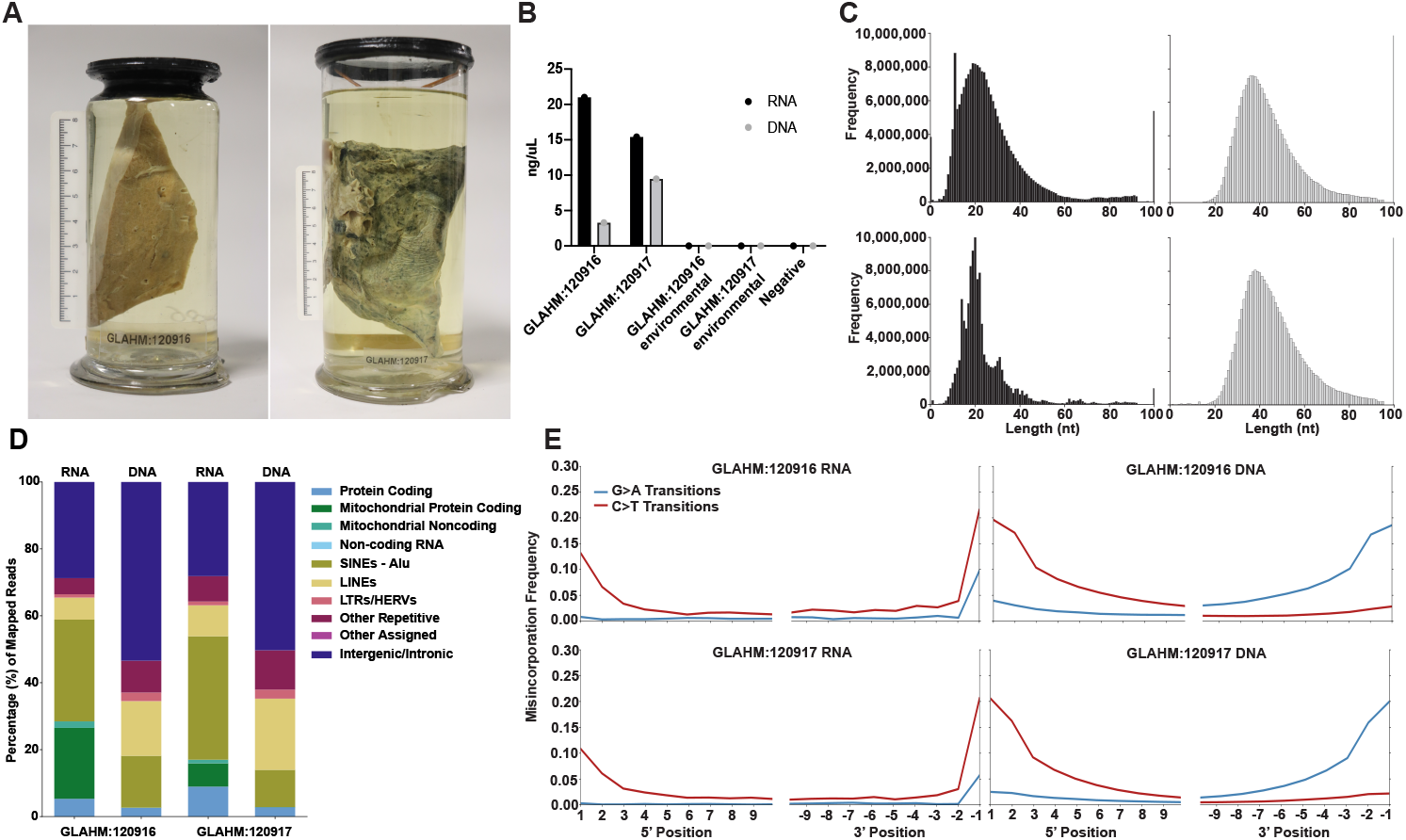
Co-isolation of aDNA/aRNA from non-formalin fixed human lung samples. (**A**) Lung specimens GLAHM:120916 (left) and GLAHM:120917 (right) preserved in alcohol solution. (**B**) Concentrations of co-isolated RNA and DNA. Measurements reported from the Qubit high sensitivity RNA assay (black) and the high sensitivity dsDNA assay (gray). 100 uL of preservation liquid was taken from each specimen and used as an environmental control. 100 uL of water was processed as a negative control. (**C)** Histograms depicting length distributions of adapter trimmed RNA and DNA libraries. RNA (black) and DNA (gray) libraries of GLAHM:120916 (top row) and GLAHM:120917 (bottom row). **(D)** Distribution of human mapped reads across biotypes. Assignments relative to the hg38 GENCODE with repetitive elements (Supplementary Materials). Assignments are labeled in color. **(E)** Misincorporation frequencies of RNA and DNA reads relative to hg38. The plot displays the C>T (red), and G>A (blue) misincorporation frequencies (y-axis) calculated using mapDamage(*29*). Misincorporation rates of the complement strand are indicated by negative values on the x-axis.

To limit spurious assignments, only reads >30nt long were mapped to the human genome, resulting in 0.03% and 0.13% endogenous RNA for GLAHM:120916 and GLAHM:120917, respectively (**Table S1 & Supplementary Materials**) (*22*). Mapped reads were subsequently classified using featureCounts (*23*). While the distribution of features among our DNA libraries closely reproduces those found in the human genome(*24, 25*), Alu retrotransposable elements and mitochondrial RNA (mtRNA) were detected at higher levels within the RNA libraries (2.53X (SINEs - Alu) and 390.52X (mtRNA) average fold enrichment) (**Fig. 1D**). Given that mitochondria’s bidirectional transcription and overlapping gene architecture can generate double stranded RNA (dsRNA) (*26*–*28*) and Alu elements are estimated to account for 80% of all endogenous dsRNA(*27*), we hypothesize that this enrichment arises from a preferential conservation or recovery of dsRNA molecules. To corroborate this, we leveraged our strand-specific RNA library preparation to identify the directionality of RNA reads mapping throughout the mitochondria. In doing this, we confirm that the mtRNA heavy and light strands are both present (**Fig. S1**).

Previous aRNA studies have reported molecular damage patterns similar, but not identical, to aDNA(*12*–*15*). However, the specific profiles of aRNA damage are still unclear and likely depend on the preservation context. To directly compare damage profiles between aDNA and aRNA in the same specimen, damage patterns for human mapped RNA and DNA were estimated using mapDamage (*29*). Analysis of the human mapped DNA shows canonical aDNA random depurination patterns (**Fig. S2A)** (*30*). Interestingly, this pattern is absent in the RNA reads (**Fig. S2B)**. It has been described that RNA has a slower rate of depurination than that of DNA and instead relies on the nucleophilic attack of the 2’OH of the ribose to the 5’ phosphate group of the downstream nucleotide, suggesting that depurination is not driving the aRNA fragmentation process (*11, 31*). Furthermore, we observe C>T transitions and base frequencies of the RNA that are distinct from the DNA (Fig. 1E). Specifically, there is a significant increase in the frequency of thymine’s at the last position of the RNA reads, corresponding to an increase in C>T transitions at the same location (**Fig. 1E & Fig. S2B)**. This may be explained by RNases specific for dsRNA, such as RNase III, that leave a 2nt overhang at the 3’ end of the dsRNA it targets, making them susceptible to random cytosine deamination (*32*). Notably, C>T misincorporation is not present in the environmental or negative controls **(Fig. S3**). Overall, these damage analyses indicate that the RNA reads contain distinct *post-mortem* associated damage.

### Reconstruction of an 18^th^ century Human Rhinovirus A genome

To characterize the aRNA virome preserved in these lung specimens, we performed both k-mer (Kraken2) and alignment-based (CZID) analyses on high-complexity non-human mapped reads (**Supplementary Materials**)(*33, 34*). Specimen GLAHM:120916 contained more viral diversity than GLAHM:120917, with the largest number of k-mers assigned to Human Rhinovirus (HRV) A, a positive-sense single-stranded RNA virus of the *Picornaviridae* family that is the most frequent causal agent of the common-cold (**Fig. 2A**)(*35*). The abundance of HRV A assigned reads in GLAHM:120916 was also identified by CZID (**Fig. S4A**). There were no reads assigned to HRV A in the negative or environmental controls. Interestingly, CZID analyses of the GLAHM:120916 aDNA libraries suggest co-infection with respiratory disease related bacteria such as *Streptococcus pneumoniae, Haemophilus Influenzae*, and *Moraxella Catarrhalis* (**Fig.S4B**).

**Fig. 2.**
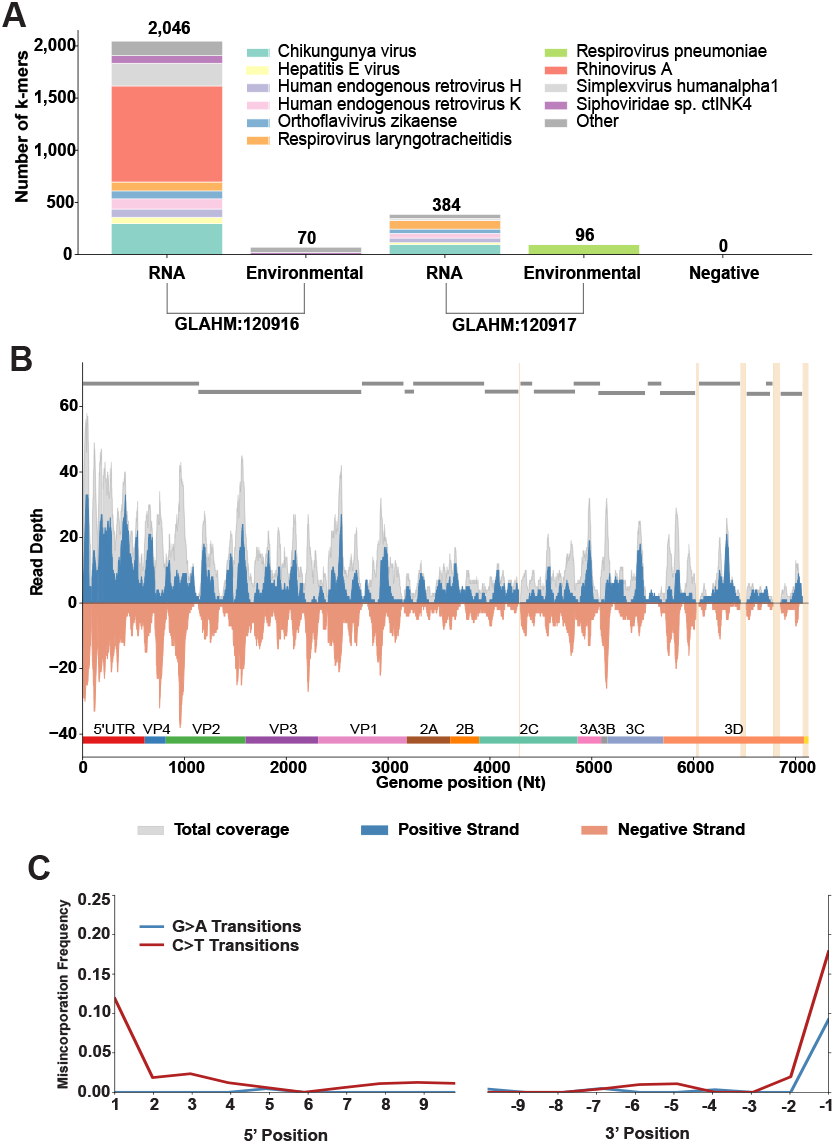
Reconstruction of an 18^th^ century human rhinovirus A genome. **(A)** Number of viral k-mers identified by Kraken2 on each lung specimen and control RNA libraries. K-mers per viral species were extracted using the Pavian suite, and all pathogens identified within the negative control library were filtered out of the subsequent libraries to remove spurious assignments. Viral species assignments are indicated by different colors and the total number of k-mers assigned per library are shown at the top of each bar. (**B)** Strand specific coverage of RNA reads mapping to the *de novo* assembled rhinovirus A genome. Positive strand coverage of the virus is displayed in blue, negative strand coverage is displayed in orange, and total coverage is represented in gray.Yellow vertical lines show gaps within the reads mapping to the assembled genome. MegaHit assembled contigs are represented by the gray bars at the top of the plot. A diagram of the HRV genome is displayed at the bottom of the plot. (**C)** Damage pattern of GLAHM:120916 RNA reads mapping to the *de novo* assembled HRV A genome. The plot displays the C>T (red), and G>A (blue) misincorporation frequencies (y-axis) calculated using mapDamage(*29*).Misincorporation rates of the complement strand are indicated by negative values on the x-axis.

To mitigate the bias associated with reference-based mapping, GLAHM:120916 RNA reads from a deeper sequencing run were *de novo* assembled into contigs using MegaHit (**Supplementary Materials**) (*36*). In total, 6,823 contigs were assembled, of which 16 contained rhinovirus sequences that were used to reconstruct ∼97% of a Rhinovirus A genome, with remaining gaps located primarily in the 3D gene and 3’ UTR (**Fig. 2B**). The concatenated GLAHM:120916 RNA libraries were then mapped back to the *de novo* assembled genome (**Supplementary Materials**). The average depth across the historical HRV A genome was 12.14X, with the highest read coverage found at the Internal Ribosomal Entry Site (IRES) in the 5’ UTR (28.38X) (**Fig. 2B**). We observe balanced coverage for both the positive and negative strands, suggesting the recovery of dsRNA replication intermediates. Interestingly, the length distribution of reads mapping to the historical genome is on average longer than the RNA reads mapping to the human genome (**Fig. S5 & Fig. S6)**. Misincorporation analyses show damage patterns similar to the human mapped aRNA, indicating that the reads mapping to the *de novo* assembled HRV A genome contain *post-mortem* associated damage (*37*) (**Fig. 2C**). Furthermore, the environmental control had no reads mapping to the *de novo* HRV A genome (**Table S2**). Collectively, our analyses indicate that the reconstructed genome represents that of an authentic Human Rhinovirus A recovered from a lung specimen preserved ∼250 years ago.

### Characterization of the Historical HRV A Genome Compared to Modern Diversity

Modern HRV sequences are extremely diverse, with 81 confirmed genotypes within HRV A (*Enterovirus alpharhino*) alone (*38, 39*). We observed that the coding sequence (CDS) of the historical HRV A genome has the highest nucleotide (86.24%) and amino acid (93.8%) identity to the ATCC reference sequence of HRV A19 (accession: FJ445119.1). Sequencing and patristic distance (*p-*distance) comparisons between rhinovirus VP1 genes are used to delineate rhinovirus genotypes from clinical data(*40*). Under current standards (>13% *p-*distance), the historical HRV A genome represents a distinct genotype from all published HRV A19 sequences (**Fig. S7 & Supplementary Materials**) (*41*). Furthermore, we observe 103 amino acid differences between the historical HRV A CDS and any HRV A19, but only four amino acids are unique to the historical HRV A compared to all HRV A genotypes across VP1, 2C, and 3C (**Fig. 3A & Supplementary Materials**). Of note, none of these unique changes falls within catalytic or structural domains. The low number of unique mutations may suggest either convergent evolution or deep historical recombination events. To further investigate these possibilities, we performed sliding-window maximum-likelihood (ML) analyses across the CDS, comparing the historical genome to representative HRV A genotypes. While these analyses show evidence of recombination among several HRV A genotypes, the historical HRV A genome is consistently immediately adjacent to HRV A19 across all windows, suggesting that HRV A19 has evolved largely in a tree-like fashion, without detectable introgression from other genotypes, for at least the past ∼250 years (**Fig. S8**).

**Fig. 3.**
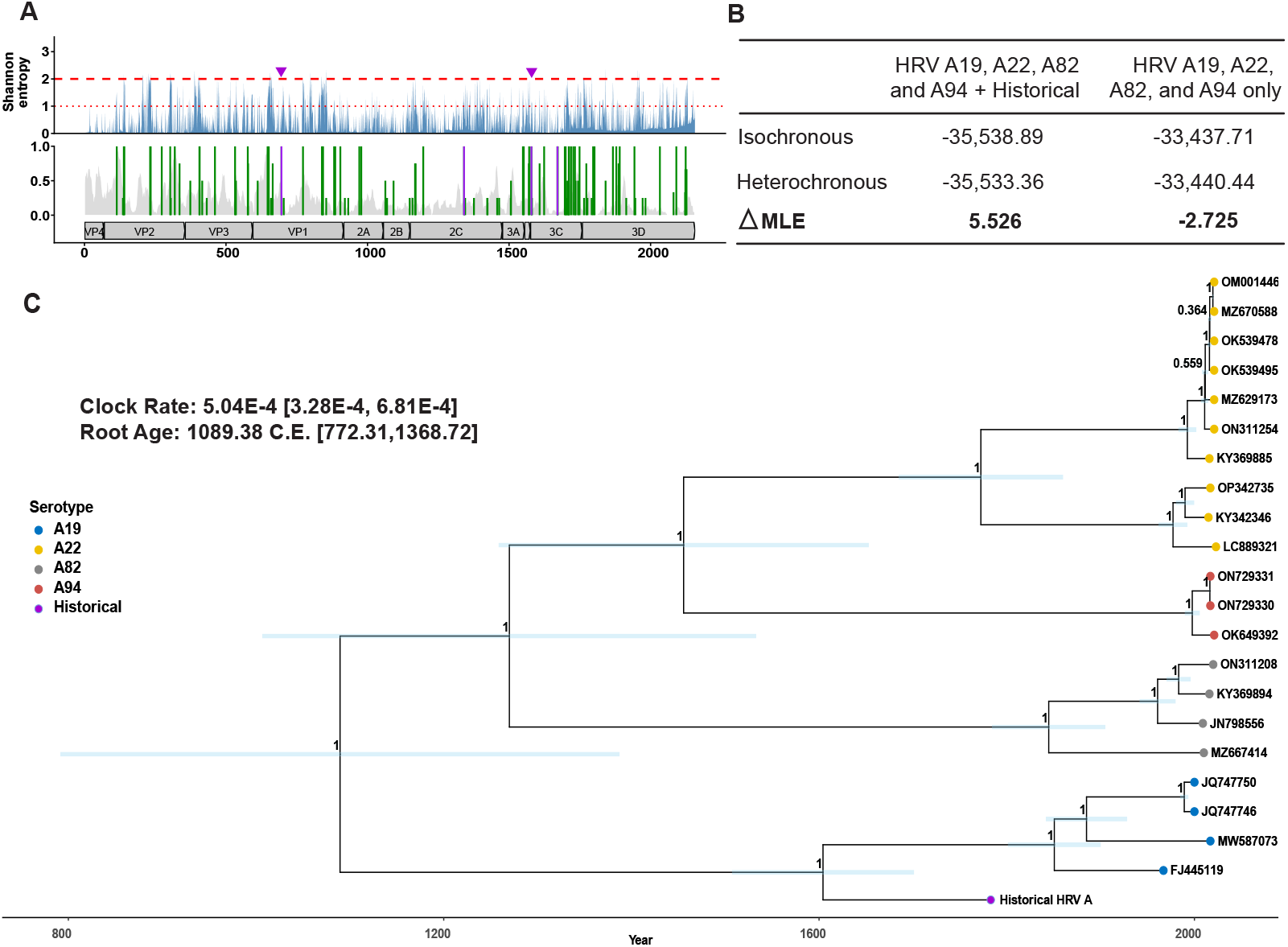
Evolutionary characterization of the 18^th^ century HRV A genome: **(A)** Amino acid (AA) variation in the historical HRV-A genome. The bottom panel shows AA differences across the CDS between the historical HRV A and any HRV A19 (green lines) and AA positions that are unique to the historical HRV A relative to all HRV A sequences (purple lines). Green line height indicates the proportion of HRV A19 sequences differing at each position. Historical HRV A genome coverage is shown in the background, with coding sequence annotations below. The top panel shows Shannon entropy per AA position (blue). Purple arrows mark the two historical-specific AAs with near-zero entropy (∼0.01). Dashed and dotted red lines denote highly (>2) and moderately (>1) variable positions, respectively. (**B)** BETS analyses displaying the log Bayes Factor (Δ*MLE*) from BEAST trees including tip dates (heterochronous) and those not including tip dates (isochronous). The Marginal Likelihood Estimates are calculated from BEAST trees utilizing a Generalized Stepping-Stone Sampling analyses. A Δ*MLE* greater than 3 is sufficient to assume strong temporal signal(44). The 95% HPD of the mean substitution rate and root age are annotated in brackets. (**C)** Time-calibrated maximum clade credibility (MCC) BEAST tree under a strict molecular clock and coalescent constant population prior utilizing full-genome sequences of genotypes A22, A82, A94 and A19 (indicated in color). The 95% HPD of each node is shown by the blue horizontal bars and the posteriors for each node are displayed as a number, 0-1. Estimated time (years) is displayed on the x-axis.

ML phylogenetic analyses placed the historical genome basal to the HRV A19 clade, and closer to the most recent common ancestor of all related genotypes (HRV A22, A82, and A94) than all modern HRV A sequences, consistent with the historical nature of our genome (**Fig. S9A**). Root-to-tip regression analyses shows temporal signal (R^2^ =0.175) (**Fig. S9B & Supplementary Materials**) (*42, 43*). A BEAST generalized-stepping stone analysis on heterochronous and isochronous trees was also preformed both in the presence and absence of the historical genome to compare the Marginal Likelihood Estimates (MLE) for temporal signal, according to the Bayesian Estimate of Temporal Signal (BETS)(*44*). BETS analyses detected strong temporal signal with the dataset that included the historical HRV A genome (Δ*MLE* = 5.52) whereas the BETS on the data without it did not (Δ*MLE* = −2.73)(**Fig. 3B**)(*45*). Notably, the inclusion of the historical HRV A genome shows a slower substitution rate of 5.04E-4 [3.286E-4, 6.808E-4] substitutions/site/year and estimates the time to most recent common ancestor (tMRCA) of this clade to 1089.38 C.E. [772.31, 1368.72] when using a strict clock (the best fitted model) with a coalescent constant population tree prior and GY94 substitution model(**Fig. 3C)**. This date predates previous estimates for the tMRCA of all HRV A genotypes, which were dated to approximately 1469 C.E. [1307.83–1574.61] and underscores a well-recognized limitation of molecular clock analyses, in which tMRCAs are often systematically underestimated due to challenges in extrapolating short-term substitution rates across deeper evolutionary timescales (*46*–*48*). As such, even the older tMRCA estimates obtained in this study with the inclusion of the historical genome are likely to represent conservative lower bounds. Nevertheless, the discovery and sequencing of the historical HRV A sequence provide better resolution to recalibrate molecular clocks previously calculated based on modern sequences alone and redefines our understanding of the evolutionary time scale of these viruses in the human population.

## DISCUSSION

Our study demonstrates that alcohol-preserved human tissues contain recoverable aRNA, enabling the reconstruction of a full RNA virus genome from a specimen nearly 250 years old— the oldest human-associated RNA virus recovered to date—indicating that RNA preservation in historical human remains is not limited to desiccated or permafrost environments but can extend to archived medical specimens even without formalin fixation. We observed an increased coverage across regions of the reconstructed historical HRV A genome predicted to form stable RNA secondary structures, including the IRES within the 5’ UTR. Such enrichment is consistent with observations of ancient ssDNA viruses like parvovirus B19, where double-stranded features (e.g., inverted terminal repeats) exhibit enhanced preservation (*49*). Consistent with this idea, we detected approximately even strand coverage across the historical HRV A genome suggesting the recovery of dsRNA intermediates, which are only present when the virus is actively replicating (**Fig. 2B**). Furthermore, we observed a high proportion of host RNAs derived from repetitive elements – including Alu sequences which have been shown to form highly structured stem-loops and are immunostimulatory – and potential mitochondrial dsRNA(*27*). This preferential preservation of dsRNA molecules has been described previously in dried entomological museum specimens (*27, 50*). Together, these observations suggest that dsRNA fragments disproportionately contributed to the surviving aRNA pool in non-formalin preserved specimens.

Our analyses show that the historical HRV A genome represents an extinct lineage closely related to HRV A19 and suggests a dynamic turnover of HRV A genotypes. The inclusion of the historical HRV A genome into time-calibrated trees showed reduced intra and inter-genotype substitution rates for HRV A (*38, 48*) (**Fig. 3C & Fig. S11**). However, these estimates might be confounded by site saturation during this temporal range or indicate that the selective pressure on HRV A was not constant across time. For example, the historical HRV A unique mutations within the structural VP1 protein (133 I) and the viral 3C protease (63 T) occur at sites that are otherwise highly conserved across modern HRV A (Shannon entropy ∼0.01), potentially representing context dependent adaptations to the particular historical environment or social behaviors (**Fig. 3A)**. Interestingly, the diversity between the historical HRV A and modern HRV A19 along the structural genes (VP4-VP1) falls in regions of moderate to high diversity (Shannon entropy >1), supporting that antigenic drift may have also contributed to the diversification from HRV A19.

The addition of a single historical genome provided strong temporal signal to reveal that the tMRCA of modern subtypes HRV A19, A22, A82, and A94 predates previous estimates of the entire diversity of HRV A when using only modern sequences(*48*). The newly inferred medieval origin is likely an underestimate of this clade yet supports the deep evolutionary relationship of HRV with human populations and suggests a substantially more ancient origin than previously appreciated of all HRV A. This interpretation is consistent with molecular dating of HRV C VP4/2 genes, which estimated their origin to ∼1,800 years ago (*51*), as well as depictions in ancient Egyptian text, where descriptions of the common cold indicate that respiratory pathogens such as HRV were circulating over 3 thousand years ago (*52*). Together, their modest basic reproductive number (R_0_=1.2-2.6), exclusive human tropism and extensive genotype diversity, suggest that HRVs were likely introduced in human populations prior to human urbanization(*53*). Our phylogenetic analyses remain limited by the narrow temporal range of available modern human rhinovirus genomes, which are predominately sampled from 2020 onward (*38*) and by the possibility that rapid lineage turnover obscures an accurate estimation of deeper divergence times. Overall, this study shows that incorporating ancient genomes can meaningfully refine deep evolutionary relationships that remain undetectable when relying solely on modern genotypes. However, the recovery of a single historical genome provides only a partial view, underscoring the need for additional ancient sequences to fully resolve the long-term evolutionary history of HRV.

Our extraction protocol uniquely enables the dual isolation of DNA and RNA from the same sample, providing a comprehensive view of the molecular history of museum specimens. In addition to the reconstructed Rhinovirus A, top hits for GLAHM:120916 include known pathogens that cause pneumonia including *Streptococcus pneumoniae, Moraxella catarrhalis*, and *Haemophilus influenzae* (**Fig. S4B**). Rhinovirus infections are typically limited to the upper respiratory tract but are known to infect the lower respiratory tract and cause severe disease when associated with bacterial co-infections (*54*). This clinical scenario is well recognized today but rarely described in 18^th^-century medical texts. William Cullen (1710–1790) offered a rare account, noting that an apparently simple catarrh might fall upon the lungs and progress to fatal *peripneumonia notha* in elderly or vulnerable patients(*55*). The clinical description of specimen GLAHM:120916 aligns closely with concurrent bacterial-viral infection that exacerbated their respiratory disease and ultimately contributed to their death. Human rhinoviruses were not identified until the mid-20^th^ century, with the first HRV isolated in 1956(*56*). In this light, our work offers a rare opportunity to connect clinical descriptions from the late 18^th^ century with the viral pathogens that would not be recognized until almost two hundred years later.

Overall, this work pushes back the known preservation window for human-associated RNA viruses and demonstrates that meaningful genomic information can survive for centuries under non-ideal archival conditions. These findings broaden the temporal and contextual range accessible for paleovirology studies and highlight museum and medical collections, such as The Hunterian, as underexplored reservoirs of historical RNA viruses. Continued exploration of these resources will be essential for illuminating the long-term evolution of RNA pathogens that have shaped human health for centuries.

## ACKNOWLEDGEMEMTS

We would like to acknowledge the individuals whose lung tissues were preserved in The Hunterian Anatomy Museum (University of Glasgow, Scotland) and included in this study. Although their identities and personal histories remain largely unknown, their lives, illnesses, and struggles are partially illuminated through this work. We hope that this study offers a measure of recognition and respect and contribute to doing justice to their stories. We extend our sincere gratitude to Jesse Bloom and David Veesler for helpful comments during the editing process. This research was approved by The Hunterian’s Collections Development Group, the Humanness Forum, and by Fred Hutchinson Cancer Center IRB (FHIRB0011112), and IRE in compliance with USG DURC/PEPP Policy. We thank Carina Díaz Uribe, Christian Molina Aguilar, Jair Santiago García and Luis Aguilar for technical support. Paleovirology work in the Blanco-Melo lab is supported by the National Institute of Allergy and Infectious Disease grant DP2AI177896, the Kinship Foundation Searle Scholars Program, and the Immunology and Vaccine Development Program at the Fred Hutchinson Cancer Center. Work in the Ávila-Arcos lab is supported by the Secretaría de Ciencia, Humanidades, Tecnología e Innovación (SECIHTI) grant CF-2023-G-957. Jon Bang Ploug was supported by the Danish National Research Foundation through the PandemiX Center of Excellence grant DNRF170.

## DATA AVAILABILITY

The reconstructed historical HRV A genome from this study is available in GenBank (accession PX830975). Reads that mapped to the historical HRV A consensus genome is available on SRA (SRR36652898). These files and all alignment, BEAST XML, and tree files are available in Dryad (DOI: 10.5061/dryad.h1893201q) and https://github.com/BlancoMeloLab/Historical_HRVA.

