## Supplementary Materials for "Recovery of an 18^th^ Century Rhinovirus Genome through Ancient RNA Isolation of Human Lungs"

### Materials and Methods

The Hunterian Anatomy Museum – Specimen Information

William Hunter (1718–1783) was one of the most influential anatomists of eighteenth-century Britain. As physician to Queen Charlotte and founder of the Great Windmill Street anatomy school in London, he developed one of the period's most extensive anatomical and pathological collections, comprising roughly 3,000 preparations(*57*). The collection provided a comparative framework for studying disease at a time when most physicians relied on isolated case histories. Matthew Baillie's *Morbid Anatomy of the Human Body* (1793), the first English-language textbook to classify diseases by organ system, drew directly on his uncle's collection, to which he had access after Hunter's death.

The specimen GLAHM:120916 (previously numbered T.86 in Hunter's original catalogue and then 18.19 in Teacher's 1900 catalogue) derives from this collection. The catalogue entry is as follows: *"A portion of lungs consolidated into a mass like the Liver, where of course the air cells were nearly obliterated: from a woman in the dissecting room" The lung retains a certain degree of the natural spongy texture; the consolidation does not appear to be pneumonic, but rather what would result from compression by pleuritic effusion, or collapse from obstruction of a bronchus. Microscopic examination gave doubtful result, but favoured the idea that the condition was pneumonia at an early stage* (*57*).The provenance note indicates that the specimen was obtained through the school's dissecting room rather than from Hunter's private clinical practice.

Specimen GLAHM:120917 (previously numbered 18.20 in Teacher's 1900 catalogue) was added to the University of Glasgow collections by Dr Allen Thomson, Regius Professor of Anatomy from 1848 to 1877. The catalogue entry is as follows: *This specimen, of which there is no history, consists of a portion of a lobe of the lung, showing a large haemorrhagic infarction. It is of a deep brown colour, of triangular shape, its base at the surface of the lung corresponding with the area of distribution of a branch of the pulmonary artery. At its apex the artery is seen laid open; it is filled with soft blood-clot, and white thrombus adhering firmly to its walls and completely obstructing it* (*57*).

Hunter preserved wet specimens in spirits of wine (alcohol) because only in this form could "the appearances of diseases' be demonstrated"(*58*). Thomson would have prepared specimens in a similar way. This method predates the introduction of formalin fixation in the late nineteenth century and has significant implications for molecular analysis: while formalin cross-links proteins and fragments nucleic acids, alcohol-based preservation does not impose the same degree of chemical damage. The Hunterian collection—bequeathed to the University of Glasgow upon Hunter's death and transferred in 1807—therefore represents a rare archive of biological material preserved before the use of formalin.

Sampling

Biopsies were taken from different parts of lung specimens GLAHM:120916 and GLAHM:120917 to maximize the probability of recovering pathogens, summing to a total of 200 mg per specimen. Approximately 20 mg of biopsied tissue were independently processed. The preservation liquid of each lung was sampled (100 uL) and utilized as an environmental control throughout the analyses.

aRNA and aDNA co-isolation and Library Preparation

Key modifications were made to the Qiagen DNeasy Blood and Tissue Kit (catalogue ID: 69504) to increase the amount of small nucleic acids recovered from the lung samples. First, the tissue was heated in Buffer ATL at 98C for 10 minutes, as described previously, to reduce the number of macromolecular crosslinks on the nucleic acids (*16*). Immediately following this step, the tissue was homogenized by 3x30 seconds of bead beating. 20 uL of Proteinase K was then added and tissues were incubated at 56C until full tissue lysis was observed. For environmental controls, only 10-minute incubations at 56C were performed. At the column binding step, an additional 3X (600 uL) of 100% ethanol was mixed with the lysed tissue prior to loading the sample into the column. This is critical to prevent the loss of smaller nucleic acids that may dissolve in the buffer solution, thereby allowing these molecules to bind to the silica membrane. The remaining protocol was carried out following the column washing steps as described by Qiagen. The final 25 uL eluate was heated on the column at 56C for one minute prior to elution. This was repeated twice with the first eluate re-loaded onto the column to increase the quantity of nucleic acids recovered. A water control was included to assess the addition of contaminants from kit buffers, known as the *Dneasy negative* throughout the analyses. RNA and DNA concentrations were read with Qubit high sensitivity RNA and dsDNA assays, respectively.

All aliquots of the same lung specimen were then pooled and concentrated using a SpeedVac DNA 130 Vacuum Concentrator System and reconstituted at 25 uL. Samples were then divided into two, with half the eluate going into RNA library preparation and the remaining eluate used for DNA library preparation. The NEBnext small RNA library kit (E7330) was used for small RNA library preparation utilizing the NEB End Repair Module (E6050) to reverse any cyclic 3' phosphates that may have accumulated during degradation as well as repair any missing 5' phosphates. This step is necessary for adaptor ligation of degraded RNA. The NEBnext small RNA library kit follows a strand specific, adapter ligation protocol that preserves the ends of the RNA molecules and maintains the ability to identify *post-mortem* damage. We prepared these as single end libraries, as the assumed degradation can be captured by one strand. The (*59*) dsDNA single end library preparation for aDNA was used with no modifications. All libraries, including the environmental controls and negative blanks, were pooled and sequenced on the Novaseq X Plus to 100M read depth/library.

To avoid modern contamination, all sample handling, nucleic acid isolation, and Illumina library preparations were performed with proper personal protective equipment (PPE) at the sterile paleogenomics facility at the National Autonomous University of Mexico's (UNAM) International Laboratory for Human Genome Research's (LIIGH).

Bioinformatic processing of Human Mapped Reads

All libraries were initially trimmed using the FastP program to remove adapters, keeping reads with a minimum length of 30nt and a minimum read quality of 30(*60*). We observe read peaks at 100nt, which corresponds to the maximum read length of the sequencing run (100 cycle flow cell), suggesting the recovery of a significant amount of RNA and DNA molecules of 100nt in length and above (**Fig. 1C**). The trimmed reads were mapped to the human genome (hg38) following the optimized bwa aln aDNA parameters for human mapping (-l 1024 -n 0.01 -o 2) (*22*), and duplicates were removed using Picard MarkDuplicates (https://gatk.broadinstitute.org/hc/en-us/articles/13832748517275-MarkDuplicates-Picard). The use of reads >30nt for the initial analyses significantly drives down the endogenous RNA content as the peak of adapter trimmed reads is <30nt for both GLAHM:120916 and GLAHM:120917 and may be further compounded through the use of aDNA mapping parameters for the RNA libraries **(Fig 1C & Table S1**). featureCounts was run on the de-duplicated RNA and DNA to assess the composition of human mapped reads utilizing an hg38 GENCODE annotation file including repetitive elements (http://labshare.cshl.edu/shares/mhammelllab/www-data/TEToolkit/GRCh38_GENCODE_rmsk_TE_plus_human_sort.gtf) (*23*). The mapDamage V2.2.3 program was used to calculate base and misincorporation frequencies on the deduplicated human mapped reads to assess the quality of the aDNA and aRNA libraries for each lung specimen (*29*). SAMtools was used to keep unmapped reads for metagenomic profiling and were processed to remove low complexity reads utilizing the BBtools suite bbduk.sh with an entropy threshold of 0.90(*61*, *62*).

Taxonomic Profiling

A comprehensive kraken2 human virus database was created utilizing the NCBI Virus Database with filters to include human infecting viruses, partial nucleotide completeness, and to exclude most SARS-CoV-2 sequences. In total, this resulted in a fasta file containing 2,643,874 independent sequences used to build the kraken2 database. All unmapped, high-complexity reads were used as kraken2 input files with a minimum hit group of 3 and the –report-minimizer-data flag to mimic krakenuniq output (*34*). The report files were then loaded into Pavian and the CSV file comparing viral k-mers per sample was downloaded(*63*). Viral families with k-mer hits in the Dneasy negative were filtered out of all other samples to reduce noise. Simultaneously, raw sequencing files for each library were loaded to the Chan Zuckerberg Identification (CZID) online platform to corroborate the Kraken2 findings. CZID uses a comprehensive microbial database containing eukaryotes, archaea, bacteria, and virus (*33*), however it performs an extremely stringent screening process including down sampling of all host filtered libraries to a maximum of 2 million reads, a stringency that may be detrimental to the detection of highly degraded ancient pathogen genomes. Utilizing the taxon heatmap function, the reads were filtered to show non-phage viruses within the sample. The DNA libraries were also processed with CZID to assess the presence of bacterial species. With the taxon heatmap function, reads were filtered to show only bacterial "known human pathogens" to reduce noise. CSV files were then downloaded with this data, normalized by the number of alignments to the NCBI nucleotide database per million reads (NTrpm) and plotted as heatmaps **(Fig. S4).**

Rhinovirus Reconstruction

To increase the read coverage of the rhinovirus, we performed an additional Novaseq X Plus run with a 25 billion read flow cell on RNA libraries for GLAHM:120916 and its environmental control. All RNA libraries for GLAHM:120916 (e.g. 1 billion read library and 100 M read library) were processed with FastP to remove adapters, keeping reads >20nt(*60*) and then concatenated to make a master library. For *de novo* assembly with MegaHit, one extra nucleotide was trimmed from the read ends to diminish the number of damaged bases incorporated into the assembled contigs using CutAdapt (*36*, *64*). MegaHit parameters were set with non-default k-mer lengths of 17, 23, 29, and 35 to accommodate the degraded nature of the reads, where the 25$^{th}$ percentile, median, and 75$^{th}$ percentile lengths were 16, 23, and 35 nt, respectively. A rhinovirus protein database was made to include all complete rhinovirus A, B, and C proteins on NCBI as of May 2025, resulting in 1,332 protein sequences indexed as a DIAMOND database(*65*). All MegaHit assembled contigs were blasted against the protein database with the diamond blastX function, resulting in 16 rhinovirus contigs. These contigs were assembled into a draft genome and verified that no stop codons existed in the polyprotein open reading frame. The libraries without the extra base trimming were aligned back to the draft genome utilizing bwa aln stringent ancient pathogen mapping (-l 32 -n 0.1) (**Fig. 2B**)(*21*). Misincorporation frequencies were calculated using mapDamage v2.2.3 (**Fig2C**). The areas of rhinovirus contigs that did not have read coverage were not included in the final historical genome sequence for phylogenetic analyses.

<u>HRV A Genomic Characterization</u>

The *p*-distance was calculated by aligning all HRV-A19 VP1 sequences with the historical HRV A genome and performing pairwise comparisons. The number of nucleotide differences between two sequences was divided by the total number of aligned positions, yielding the pairwise *p*-distance (*41*). All modern HRV A19 was accurately called as the same genotype *(p*-distance <0.13), yet the historical HRV A was not with a *p*-distance >0.13 in all comparisons, confirming the historical HRV A is a distinct genotype (**Fig. S7**). Next, the HRV A amino acid (AA) sequences across HRV A19, including the translated historical HRV A sequence, were aligned with MAFFT and inspected in AliView v1.28 (*66*, *67*). In total, 103 AA positions in at least one HRV A19 differed from the historical HRV A. Shannon entropy analyses, which quantifies AA uncertainty at each position, indicated that 64.14% of these sites were moderately variable (entropy>1) and 4.85% were highly variable (entropy>2) **(Fig 3A)** according to (*68*). Extending this analysis to all AA sequences for HRV A identified only 4 AA unique to the historical HRV A across the VP1 (pos. 133I), 2C (pos. 245T), and 3C (pos. 63T and 185A) genes. To assess the potential of recombination with the historical HRV A genome, an alignment file containing all prototype HRV A coding sequences was split into 14 equal length alignments (471 nt). Maximum Likelihood (ML) trees were inferred for all alignments under a GTR+G substitution model with 1,000 bootstraps. The trees were then plotted as a tangle gram to assess the recombination of HRV sequences (**Fig. S8**).

<u>Phylogenetic Estimate of HRV A19 Divergence</u>

In total, only 6 HRV A19 sequences were included in our phylogenetic analyses due to the limited availability of full-length HRV genomes **(Table S3**). Sequences were aligned with MAFFT, and alignment errors were manually corrected utilizing AliView v1.28 (*66*, *67*). ML trees were inferred with IQTree2 utilizing the Model Finder parameter and 1,000 bootstraps (*69*) and temporal signal was assessed with TempEst (*42*). ML phylogenetic analyses placed the historical HRV A genome basal to the HRV A19 clade and is closer to its immediate ancestral node than to any modern HRV A19 sequence, further supporting its ancestral nature (**Fig. S10A**). Given that root-to-tip regression analyses show temporal signal ($R^2 = 0.847$), we reconstructed a time-calibrated tree using BEAST v10.5 under a Goldman-Yang 1994 (GY94) substitution model, strict molecular clock, and coalescent constant population tree prior utilizing a HRV A22 sequence as an outgroup (**Table S4**, **Fig. S10B & Fig. S11**) (*42*). The Bayesian Estimate of Temporal Signal (BETS) was performed utilizing Generalized Stepping-Stone sampling (GSS) of alignments containing the historical HRV A with (heterochronous) and without (isochronous) tip dates under the same tree parameters and a 1,000,000 MCMC chain length(*44*). The output Marginal Likelihood Estimate (MLE) of the heterochronous data was then subtracted from the MLE of the isochronous data to assess the temporality both with and without the historical genome (*44*). The heterochronous data containing the historical genome had strong temporal signal ($\Delta MLE = 12.8$) (*45*), and estimated the substitution rate of the extended HRV A19 clade to 8.00E-4 [5.39E-4, 1.07E-3] substitutions/site/year (**Table S4 & Fig S11**). Additionally, the most recent common ancestor (MRCA) of the historical HRV A and modern HRV A19 was estimated to 1670 C.E. [1580.34-1732.74] (**Fig. S11**). All BEAST log files were assessed utilizing Tracer v1.7.2 to confirm sufficient effective samples sizes (ESS >200) and the sample trees were summarized as a Maximum Clade Credibility tree with TreeAnnotator v2.7.7. All trees were subsequently viewed in FigTree v1.4.4.

Phylogenetic Estimate of Inter-Genotype Divergence

All 22 complete sequences of A19, A22, A82, and A94 were aligned for inter-genotype phylogenetic estimates utilizing MAFFT and erroneous alignments were manually corrected with AliView v1.28 (**Table S6**) (*66*, *67*)(Table S6). Using the same approach applied to the HRV A19 ML tree, we again confirmed that the historical HRV A genome falls basally to the HRV A19 clade (**Fig. S9**). Although the root-to-tip regression showed a positive $R^2$ value, the residuals were high (**Fig. 3A**), indicating substantial scatter and making the choice of an appropriate molecular clock model for Bayesian inference uncertain. To resolve this uncertainty, we performed GSS sampling analyses (100 steps) under three alternative molecular clock models—strict clock, fixed local clock, and lognormal uncorrelated relaxed clock—each paired with a coalescent constant-population prior and the GY94 substitution model. The resulting MLEs indicated that the strict clock provided the best fit model ($MLE_{strict}$= -35,533.36, $MLE_{fixed\ local}$= -35,533.66 $MLE_{uncorrelated\ relaxed}$= -35,535.29), as reflected by its lowest marginal likelihood. Accordingly, the time-calibrated analysis employed a strict molecular clock with a GY94 model and a coalescent constant-population tree prior. Temporal signal was then evaluated using BETS, following the same procedure described for the HRV A19 analyses. Posterior trees were summarized with TreeAnnotator v2.7.7 and the resulting MCC tree was visualized with FigTree v1.4.4. The mean clock rate and time to most recent common ancestor with their respective 95% highest posterior density (HPD) intervals were reported.

Human mapped DNA contaminants

When analyzing the human-mapped DNA, we detected *post-mortem* damage patterns consistent with aDNA in both our Dneasy negative and environmental controls, despite their read counts being substantially lower than the real specimens (**Table S1 and Fig. S12**). This unfortunately

suggests low-level aDNA contamination across samples in the dataset. We suspect that the source of this spurious damaged DNA is index hopping during the NovaSeq X Plus sequencing runs. Our libraries were prepared using single-index barcodes only 6nt in length, which increases the likelihood of index misassignment; using dual unique indexes in future work would enable the removal of unexpected index combinations. It is also likely that the highly degraded nature of our samples generated elevated levels of free adapters in the final libraries, which are known to further increase index hopping. Additionally, the patterned flow cell architecture of the NovaSeq X Plus has been reported to exhibit higher rates of index hopping, and the low complexity of our sequencing pools—containing only a 1% PhiX spike-in—may have exacerbated the issue.
All the information above can be found at:
https://www.illumina.com/techniques/sequencing/ngs-library-prep/multiplexing/index-hopping.html

Sequence Upload

All sequencing files uploaded to Dryad (DOI: 10.5061/dryad.h1893201q) and SRA (SRXXXX) only contain reads that were used to construct the historical HRV A genome and do not contain human reads (in accordance with the Fred Hutchinson Cancer Center IRB: FHIRB0011112).
