## Supplementary material for "Recovery of an 18^th^ Century Rhinovirus Genome through Ancient RNA Isolation of Human Lungs": Fig. S1

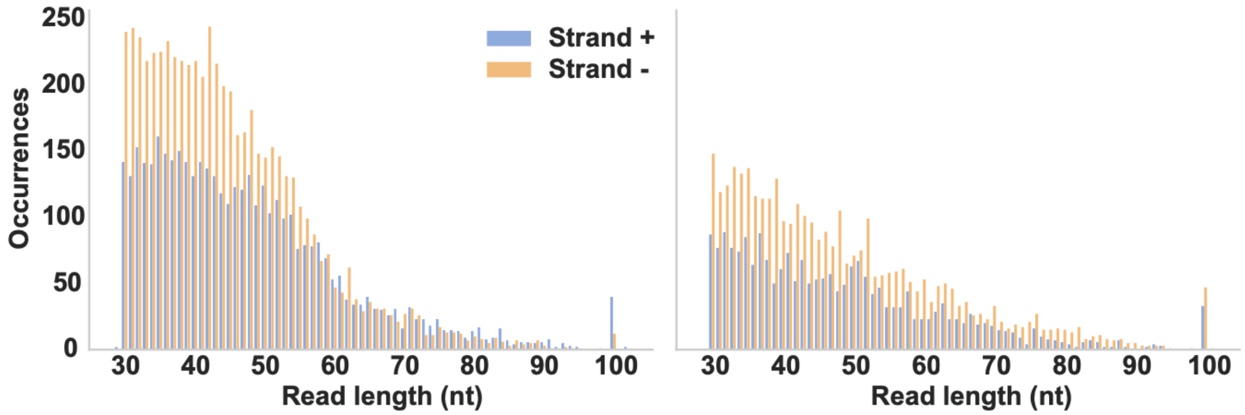


**Fig. S1. Stand specific mitochondrial RNA reads >30nt.** Length distribution of RNA reads mapping to the mitochondria. Heavy chain and light chain are represented by blue and orange bars, respectively. The mean lengths for GLAHM:120916 and GLAHM:120917 are 46.1 and 48.3nt, respectively.
