## Supplementary material for "Recovery of an 18^th^ Century Rhinovirus Genome through Ancient RNA Isolation of Human Lungs": Fig. S2

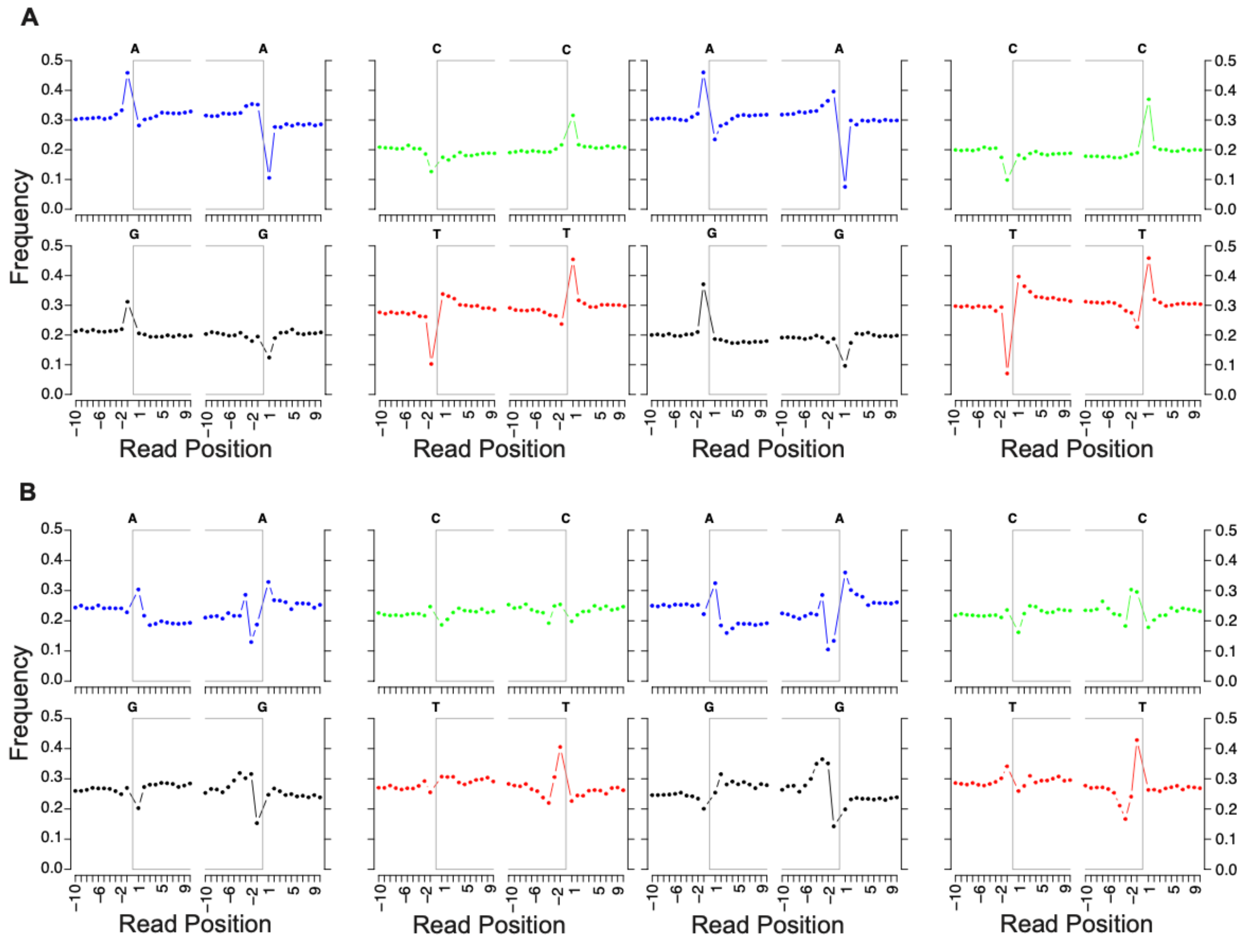


**Fig. S2. MapDamage base frequency results for RNA libraries mapping to the human genome (hg38).** The base frequency (y-axis) of adenine (blue), guanine (black), cytosine (green), and thymine/uracil (red) per position (x-axis) of mapped reads are represented within the boxed regions. Base frequencies outside the boxed region characterize the locations on the human reference adjacent to the mapped read. Positions that are negative indicate the frequencies associated with the complement strand. (**A**) Base frequency of total DNA mapped to the human genome for GLAHM:120916 (left) and GLAHM:120917(right). (**B**) Base frequency of total RNA mapped to the human genome for GLAHM:120916 (left) and GLAHM:120917(right).
