## Supplementary material for "Recovery of an 18^th^ Century Rhinovirus Genome through Ancient RNA Isolation of Human Lungs": Fig. S3

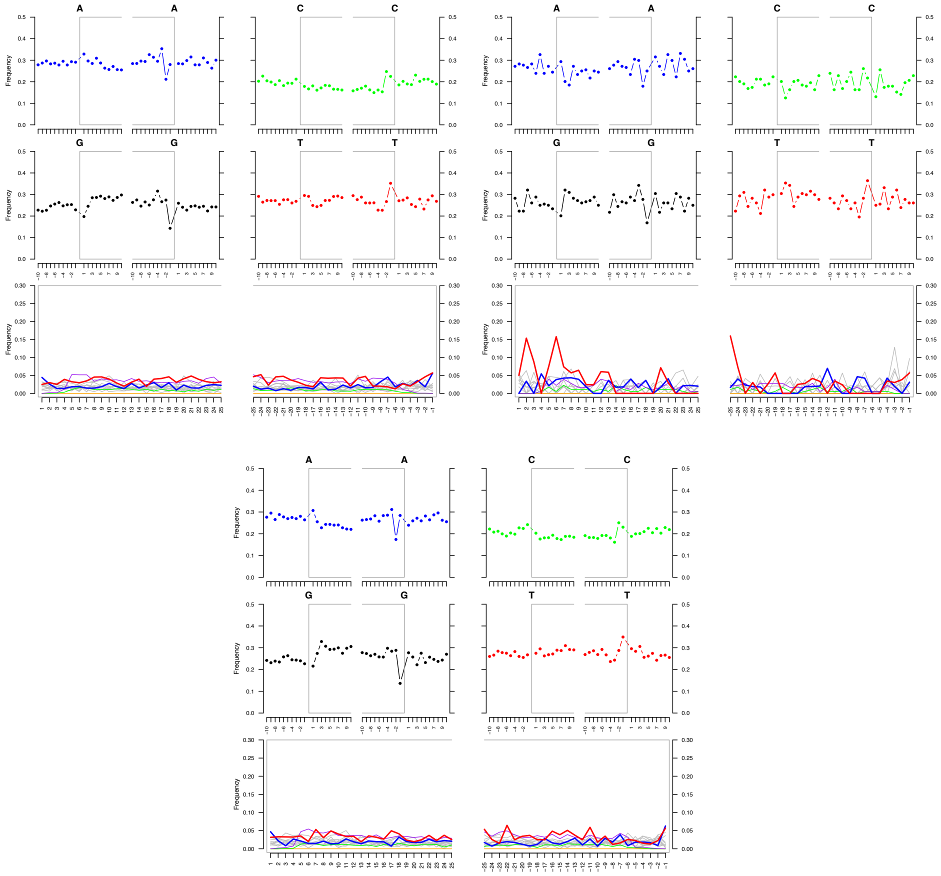


**Fig. S3. MapDamage results for environmental control and negative RNA libraries mapping to the human genome (hg38).** The top four plots are base frequency plots. The base frequency (y-axis) of adenine (blue), guanine (black), cytosine (green), and thymine/uracil (red) per position (x-axis) of mapped reads are represented within the boxed regions. Base frequencies outside the boxed region characterize the locations on the human reference adjacent to the mapped read. Positions that are negative indicate the frequencies associated with the complement strand. The bottom plot displays the C>T (red), and G>A (blue) misincorporation of reads compared to the reference. From left to right: GLAHM:120916 environmental control total human mapped RNA, GLAHM:120917 environmental control total human mapped RNA, and negative control total human mapped RNA (bottom).
