## Supplementary material for "Recovery of an 18^th^ Century Rhinovirus Genome through Ancient RNA Isolation of Human Lungs": Fig. S4

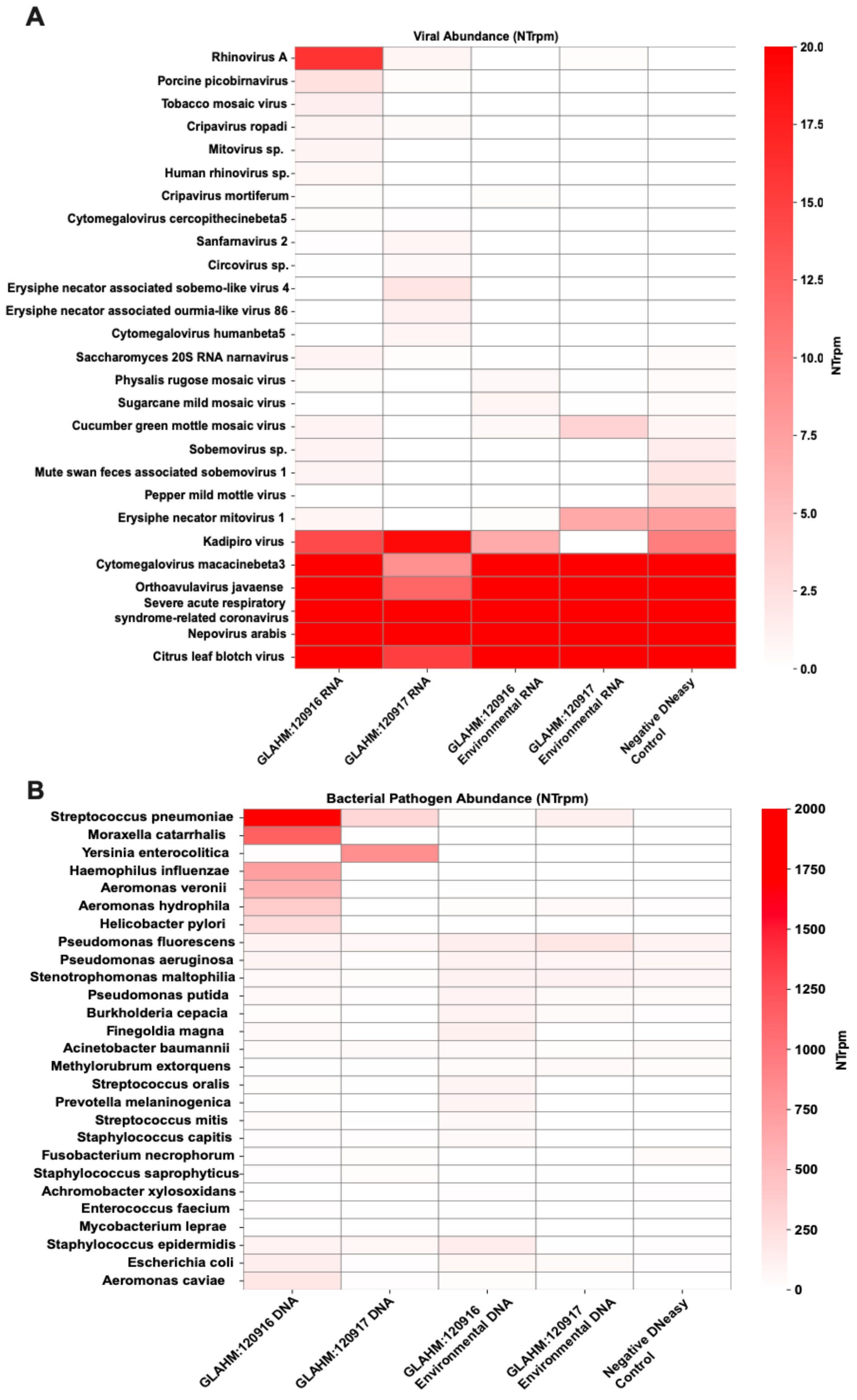


**Fig. S4. CZID taxonomic profiling of known pathogens**. (A) Distribution of viral species. (B) Distribution of known bacterial pathogens. CZID read counts are normalized as the number of reads that aligned to the nucleotide (NT) database on NCBI per million read (NTrpm). Known species identified by CZID are displayed on the y-axis. GLAHM:120916, GLAHM:120917, and their environmental controls are denoted on the x-axis.
