## Supplementary material for "Recovery of an 18^th^ Century Rhinovirus Genome through Ancient RNA Isolation of Human Lungs": Fig. S5

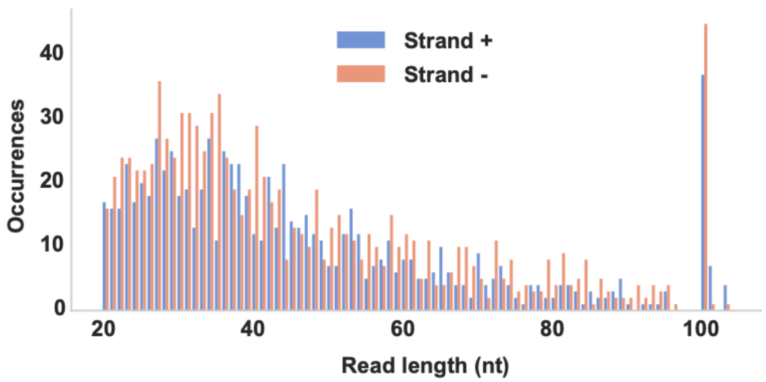


**Fig. S5. Strand specific length distribution of RNA reads mapping to the *de novo* assembled rhinovirus A genome.** Reads mapping to the positive and negative strand are shown in blue and orange, respectively. The mean length is 46.42 nt when trimmed to include reads >20nt.
