## Supplementary material for "Recovery of an 18^th^ Century Rhinovirus Genome through Ancient RNA Isolation of Human Lungs": Fig. S6

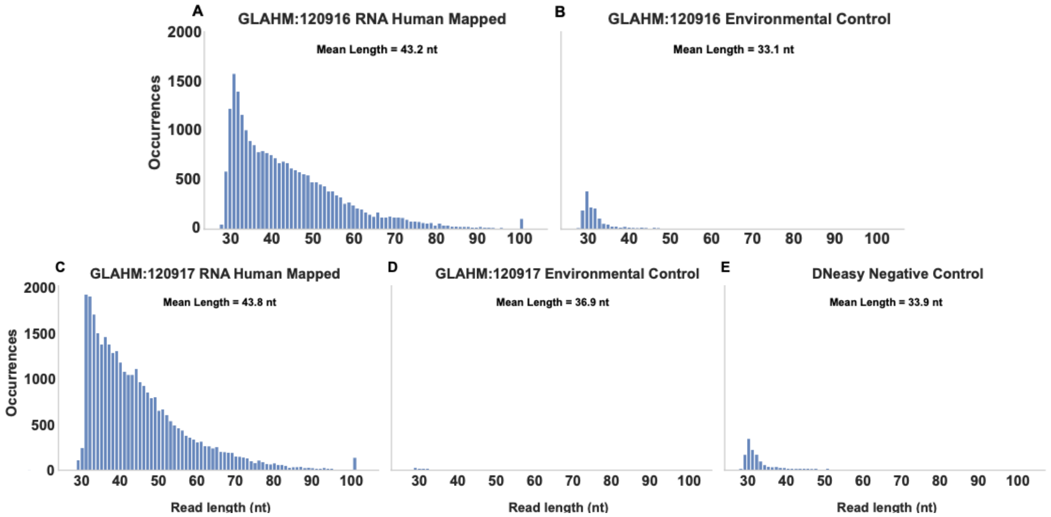


**Fig. S6. Length distribution of >30nt trimmed RNA reads mapping to the human genome (hg38)**. Bars represent the number of occurrences of each read length per RNA library. (A) GLAHM:120916 total human mapped RNA (mean length = 43.2 nt), (B) GLAHM:120916 environmental control total human mapped RNA (mean length = 33.1 nt), (C) GLAHM:120917 total human mapped RNA (mean length = 43.8 nt), (D) GLAHM:120917 environmental control total human mapped RNA (mean length = 36.9 nt), (E) Negative control total human mapped RNA (mean length = 33.9 nt).
