## Supplementary material for "Recovery of an 18^th^ Century Rhinovirus Genome through Ancient RNA Isolation of Human Lungs": Fig. S7

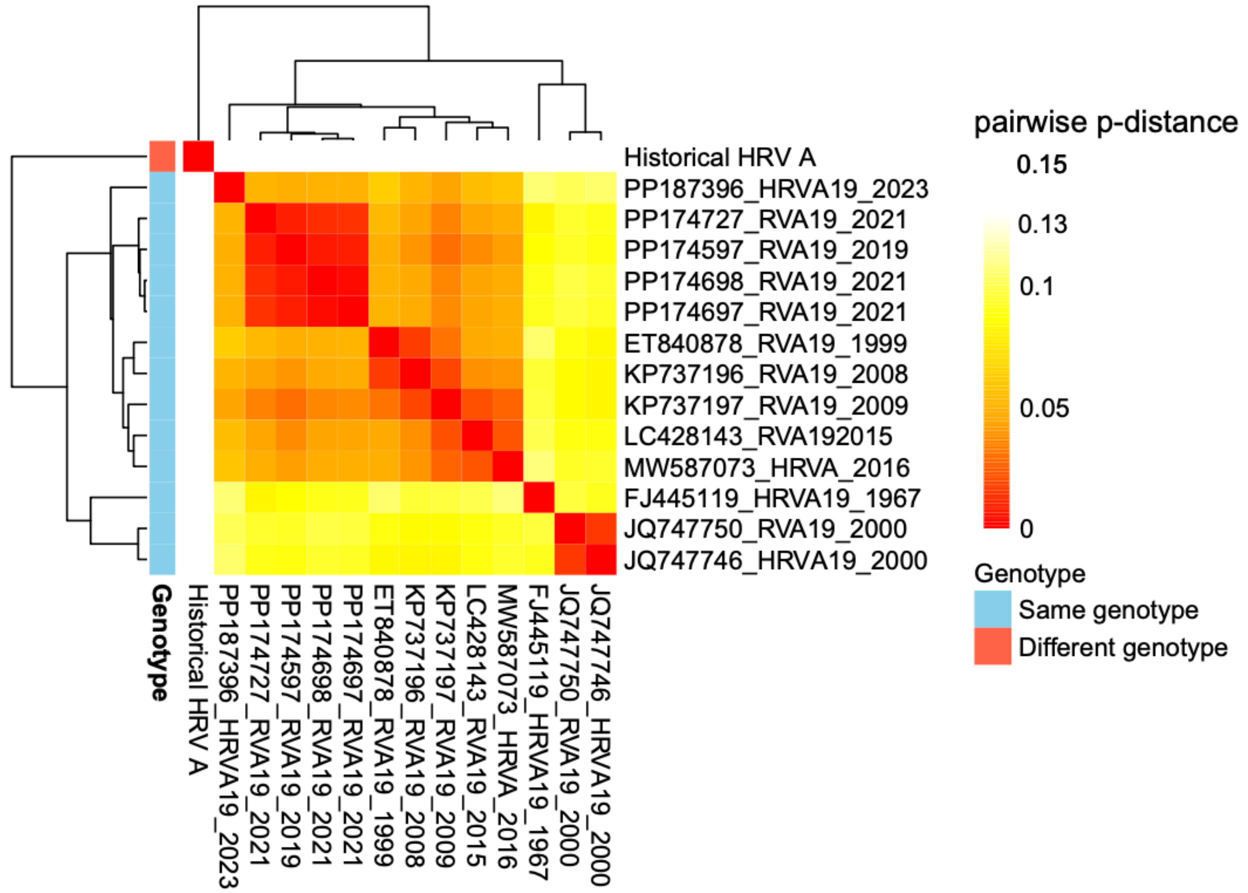


**Fig. S7. Heatmap of *p*-distances of all A19 VP1 sequences and the historical HRV VP1.** A pairwise *p*-distance was calculated between every sequence within the matrix. *P*-distances that are colored indicate two sequences are the same genotype (*p-distance* < *0.13)*. *P*-distances in white depict sequences of different genotypes (*p-distance* > 0.13).
