## Supplementary material for "Recovery of an 18^th^ Century Rhinovirus Genome through Ancient RNA Isolation of Human Lungs": Fig. S8

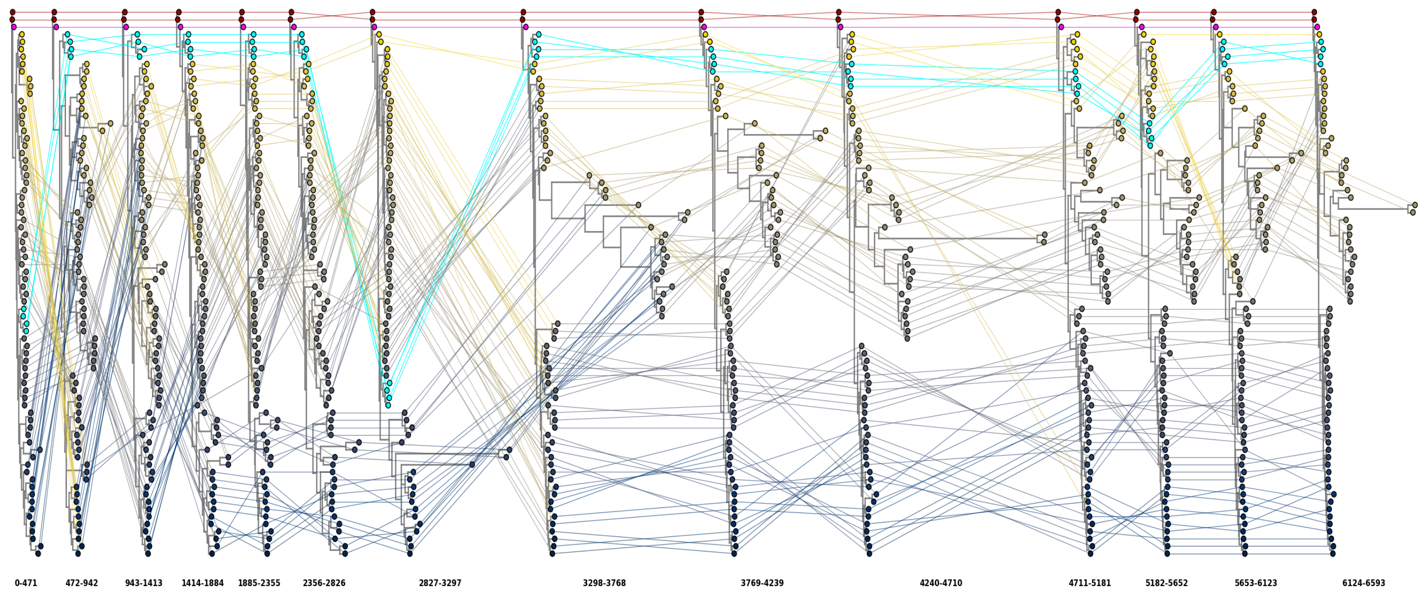


**Fig. S8. Tanglegram of sliding window ML trees across prototype HRV A coding sequences.** 14 ML trees were constructed under a GTR+G nucleotide substitution model and 1000 bootstraps. The pink tip label represent the historical rhinovirus genome, the dark red tip labels represent two HRV A19 sequences, and the blue tip labels represent the HRV A22, A82, A94, and A64 sequences. Each tip is connected by a line throughout all 14 trees to trace instances of recombination in different areas of the rhinovirus coding sequence. All trees are rooted in the A19 branch.
