## Supplementary material for "Recovery of an 18^th^ Century Rhinovirus Genome through Ancient RNA Isolation of Human Lungs": Fig. S9

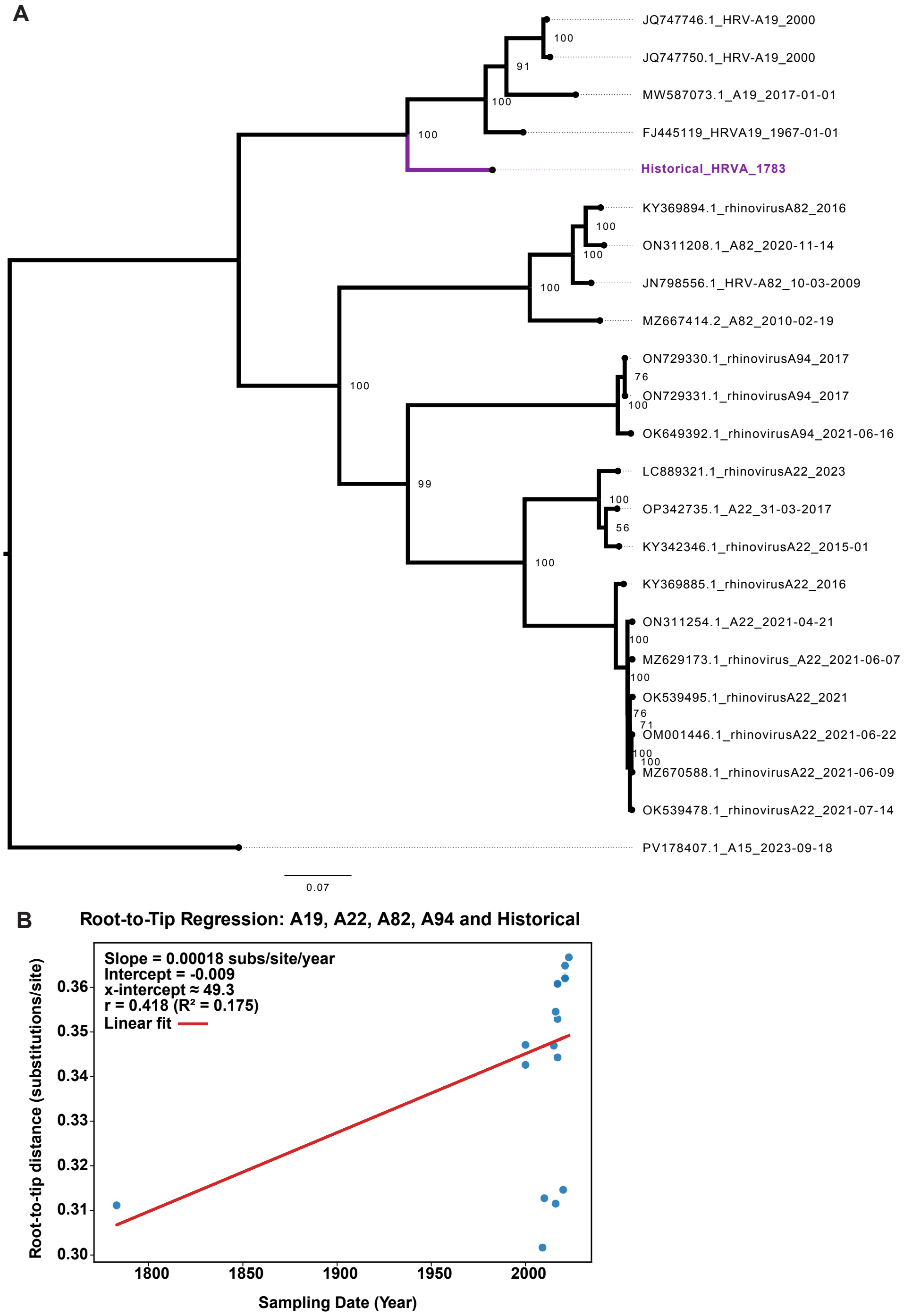


**Fig. S9. Maximum Likelihood (ML) Analyses of HRV A19, A22, A82, and A94 Clade (A)** ML Tree of HRV A19, A22, A82, and A94 and the historical HRV A genome. Historical HRV A genome is depicted in purple. A sequence of HRV A15 was used as an outgroup to root the tree. The ML tree was constructed with IQTree2 utilizing the model finder parameter (-m MFP) and preformed 1000 bootstraps. **(B)** Root-to-tip distance and sampling dates from TempEst best fitting tree. Blue dots represent genomes from corresponding ML tree (Fig. S11) plotted based on their root-to-tip distance (y-axis) and their isolation dates (x-axis). A linear regression (red line) was preformed to assess positive temporal signal ($R^2$ value > 0) and to estimate the substitution rate (slope of the line) and the age of the tree root (x-intercept value).
