## Supplementary material for "Recovery of an 18^th^ Century Rhinovirus Genome through Ancient RNA Isolation of Human Lungs": Fig. S10

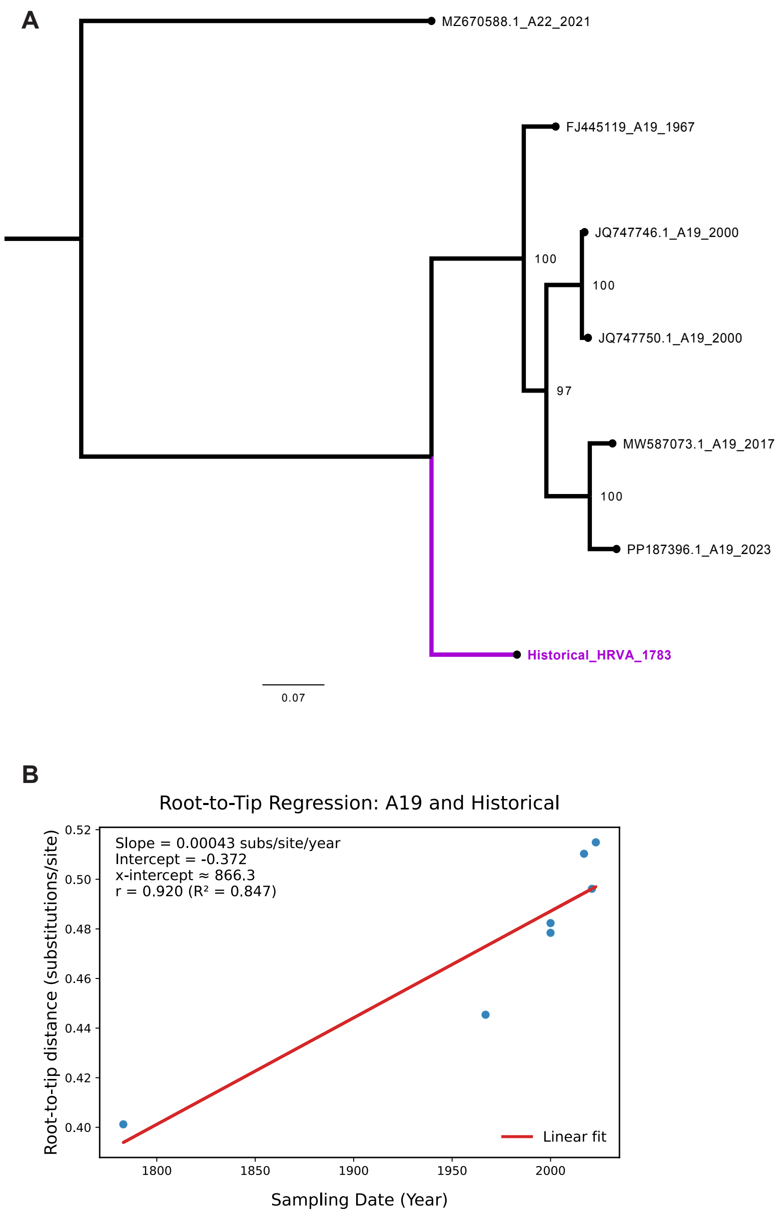


**Fig. S10. ML Analyses of the HRV A19 Clade**. (**A**) ML Tree of all HRV A19 complete genomes, and the historical HRV A genome. A HRV A22 sequence was used as an outgroup to root the tree. The ML tree was constructed with IQTree2 utilizing the model finder parameter (-m MFP) and preformed 1000 bootstraps. (**B**) Root-to-tip distance and sampling dates from TempEst best fitting tree. Blue dots represent genomes from corresponding ML tree plotted based on their root-to-tip distance (y-axis) and their isolation dates (x-axis). A linear regression (red linear fit line) was preformed to assess positive temporal signal ($R^2$ value > 0) and to estimate the substitution rate (slope of the line) and the age of the tree root (x-intercept value).
