## Supplementary material for "Recovery of an 18^th^ Century Rhinovirus Genome through Ancient RNA Isolation of Human Lungs": Fig. S11

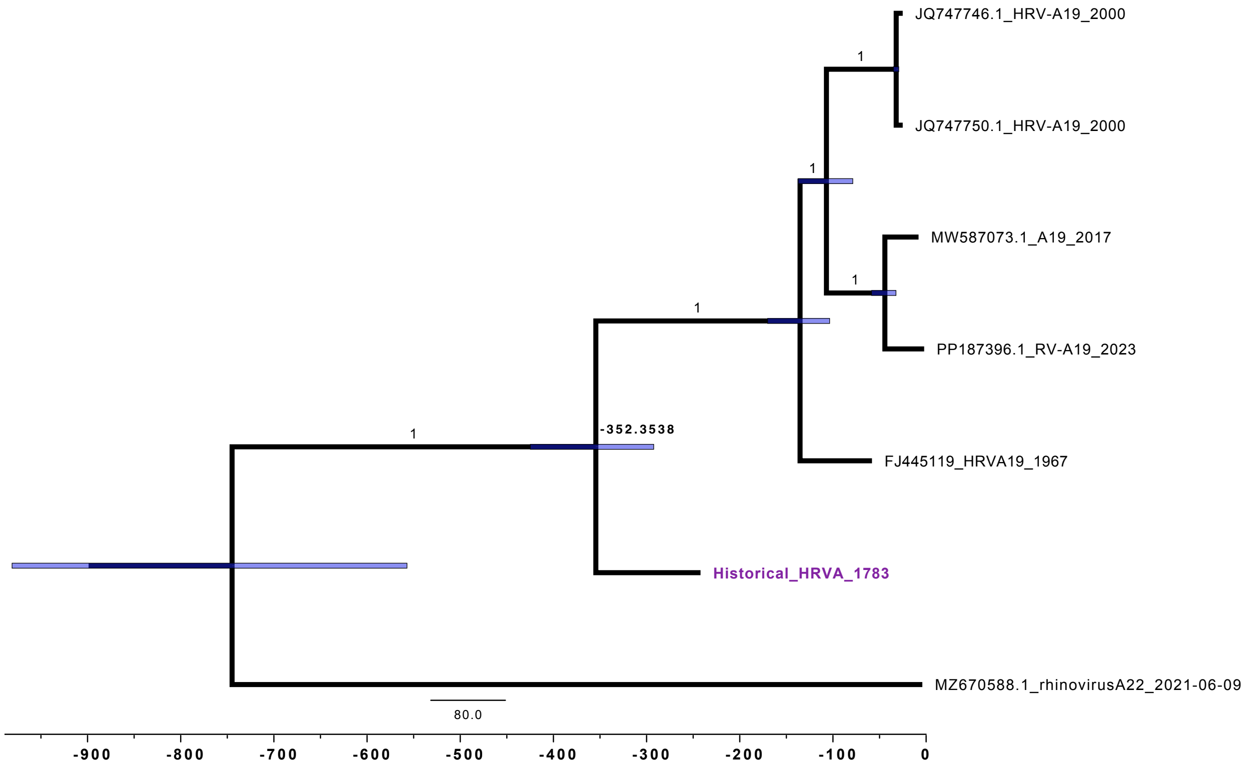


**Fig. S11. Time-Calibrated Maximum Clade Credibility Tree (MCCT) of HRV A19 clade.** The historical HRV A genome is indicated in purple. A HRV A22 sequence is included for comparison. The posterior for each node is 1, indicating strong confidence of the MCCT topology at these branch points. The 95% Highest Posterior Density of node ages are displayed by the blue horizontal bars. The scale across the x-axis represents the year in respect to the youngest sample, which was collected in 2023. This tree was constructed under a GY94 substitution model assuming a strict molecular clock and coalescent constant population size.
