## Supplementary material for "Recovery of an 18^th^ Century Rhinovirus Genome through Ancient RNA Isolation of Human Lungs": Fig. S12

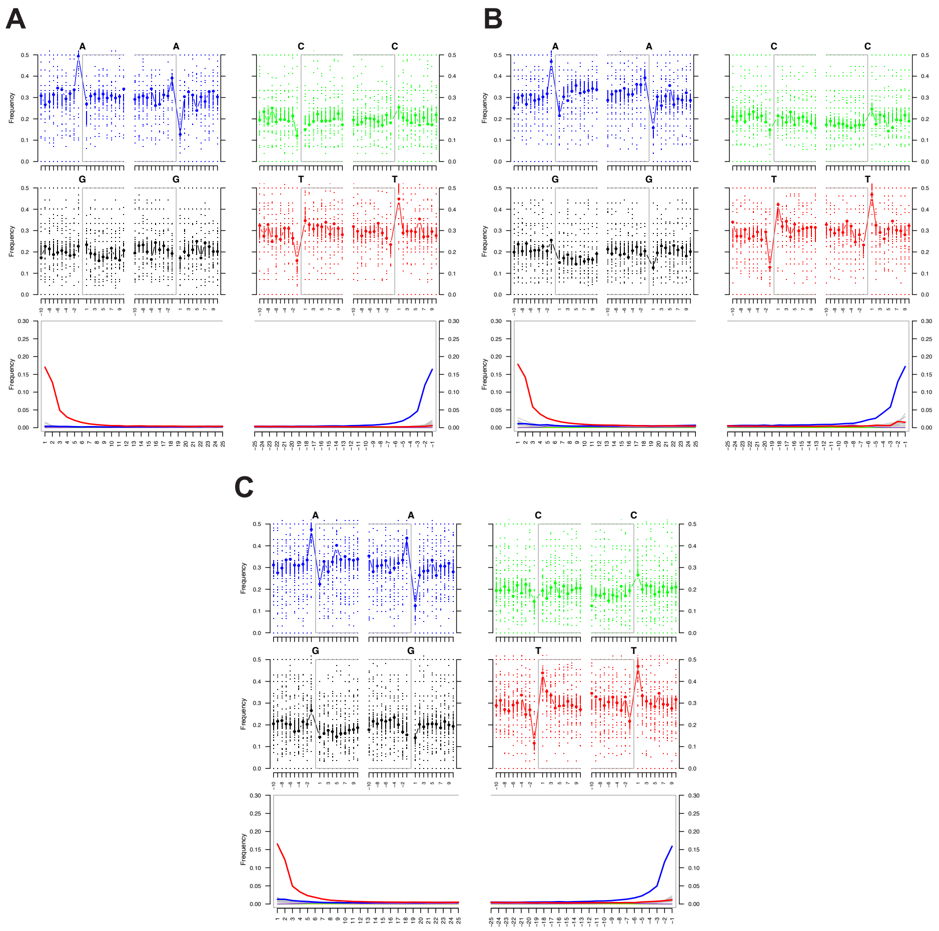


**Fig. S12. mapDamage results of DNA specimens and controls mapped to the human genome (hg38).** Human mapped reads show damage across all samples within the DNA data. (A) results for DNA mapped from specimen GLAHM:120916 environmental control, (B) results for DNA mapped from specimen GLAHM:120917 environmental control, (C) results for DNA mapped to the Dneasy negative.
