## Supplementary material for "Recovery of an 18^th^ Century Rhinovirus Genome through Ancient RNA Isolation of Human Lungs": Table S1

| Library | Molecule | Library Type | After trimming (>30 nt) | Human mapped | Reads after deduplication |
|---|---|---|---|---|---|
| 120916_R | RNA | GLAHM:120916 | 80,881,450 | 106,259 | 27,182 |
| 120916_env_R | RNA | Environmental Control | 33,225,721 | 13,053 | 1,565 |
| 120917_R | RNA | GLAHM:120917 | 26,026,347 | 63,746 | 33,535 |
| 120917_env_R | RNA | Environmental Control | 8,036,800 | 578 | 184 |
| Neg_Dneasy_R | RNA | Negative Control | 47,228,283 | 13,272 | 1,489 |
| 120916_D | DNA | GLAHM:120916 | 155,120,944 | 15,396,664 | 13,884,439 |
| 120916_env_D | DNA | Environmental Control | 67,809,830 | 232,838 | 213,680 |
| 120917_D | DNA | GLAHM:120917 | 217,710,188 | 34,331,768 | 34,302,134 |
| 120917_env_D | DNA | Environmental Control | 56,538,118 | 232,416 | 204,076 |
| Neg_Dneasy_D | DNA | Negative Control | 86,439,758 | 341,644 | 336,832 |

**Table S1. Mapping to Human Genome.** Each library was sequenced at a 100 million read depth. Adapter removed reads greater than 30nt were retained for mapping to the human genome (hg38) following the standardized human mapping bwa aln parameters described in (*22*). Deduplication was conducted with Picard Suite's MarkDuplicates.
