## Supplementary material for "Recovery of an 18^th^ Century Rhinovirus Genome through Ancient RNA Isolation of Human Lungs": Table S2

| Library | Trimmed >20 nt | Number of reads | Covered bases | Coverage | Mean Depth |
|---|---|---|---|---|---|
| Specimen | 1,300,155,682 | 1825 | 6903 | 96.8163 | 11.8663 |
| environmental control | 174,903,390 | 0 | 0 | 0 | 0 |

**Table S2. Coverage table of reads mapping to historical HRV A genome.** Specimen consists of all RNA libraries for GLAHM:120916 concatenated, to enable the full reconstruction of the historical rhinovirus. Both the specimen and environmental libraries were trimmed at 20nt to capture small reads mapping to the assembled rhinovirus genome with strict ancient pathogen mapping parameters (bwa aln -l 32 -n 0.1)(*21*). The table is the result of SAMtools coverage output(*62*).
