## Supplementary material for "Recovery of an 18^th^ Century Rhinovirus Genome through Ancient RNA Isolation of Human Lungs": Table S3

| Accession | Date | Genotype |
|---|---|---|
| JQ747746.1 | 2000 | HRV A19 |
| JQ747750.1 | 2000 | HRV A19 |
| MW587073.1 | 2017 | HRV A19 |
| FJ445119.1 | 1967 | HRV A19 |
| PV178407.1 | 2023.0465 | HRV A19 |
| Historical HRV A | 1783 | Novel |

**Table S3.** Accession numbers of HRV A19 sequences included in the HRV A19 clade phylogenetic analyses. Collection date is in decimal year format and classified genotype is listed.
