## Supplementary material for "Recovery of an 18^th^ Century Rhinovirus Genome through Ancient RNA Isolation of Human Lungs": Table S4

| | HRV A19 + Historical | HRV A19 only |
|---|---|---|
| Isochronous | -20827.8 | -18625.2 |
| Heterochronous | -20815.6 | -18624.9 |
| **ΔMLE** | **12.18** | **0.31** |
| Clock Rate [95% HPD] | 8.00E-4 [5.39E-4,1.07E-3] | 1.49E-3 [9.12E-4, 2.06E-3] |

**Table S4. BETS analysis for Temporal Signal of extended HRV A19 Clade and Outgroup**. BETS analyses displaying the log Bayes Factor ($\Delta MLE$) calculated from a generalized stepping-stone (GSS) analysis with a matching coalescent tree prior. The GSS analysis samples from tree data including tip dates (heterochronous) and those not including tip dates (isochronous). The $\Delta MLE$ was calculated for the HRV A19 clade containing the historical genome as well as the HRV A19 clade alone. A $\Delta MLE$ greater than 3 is sufficient to assume strong temporal signal(*45*). The estimated substitution rates from the MCCT are also displayed for each tree with their 95% HPD in brackets.
