## Supplementary material for "Recovery of an 18^th^ Century Rhinovirus Genome through Ancient RNA Isolation of Human Lungs": Table S5

| Accession | Date | Genotype |
|---|---|---|
| KY369885.1 | 2016 | HRV A22 |
| ON311254.1 | 2021.0548 | HRV A22 |
| MZ629173.1 | 2021.0164 | HRV A22 |
| OK539495.1 | 2021 | HRV A22 |
| OK539478.1 | 2021.0356 | HRV A22 |
| OM001446.1 | 2021.0575 | HRV A22 |
| MZ670588.1 | 2021.0219 | HRV A22 |
| LC889321.1 | 2023 | HRV A22 |
| OP342735.1 | 2017.0822 | HRV A22 |
| KY342346.1 | 2015 | HRV A22 |
| KY369894.1 | 2016 | HRV A82 |
| JN798556.1 | 2009.0247 | HRV A82 |
| MZ667414.2 | 2010.0493 | HRV A82 |
| ON311208.1 | 2020.0355 | HRV A82 |
| ON729330.1 | 2017 | HRV A94 |
| ON729331.1 | 2017 | HRV A94 |
| OK649392.1 | 2021.0411 | HRV A94 |
| JQ747746.1 | 2000 | HRV A19 |
| JQ747750.1 | 2000 | HRV A19 |
| MW587073.1 | 2017 | HRV A19 |
| FJ445119.1 | 1967 | HRV A19 |
| PV178407.1 | 2023.0465 | HRV A19 |
| Historical HRV A | 1783 | Novel |

**Table S5.** Accession numbers of HRV sequences included in inter-genotype phylogenetic analyses. Collection date is in decimal years and classified genotype listed.
